# Rapid SARS-CoV-2 whole genome sequencing for informed public health decision making in the Netherlands

**DOI:** 10.1101/2020.04.21.050633

**Authors:** Bas B. Oude Munnink, David F. Nieuwenhuijse, Mart Stein, Áine O’Toole, Manon Haverkate, Madelief Mollers, Sandra K. Kamga, Claudia Schapendonk, Mark Pronk, Pascal Lexmond, Anne van der Linden, Theo Bestebroer, Irina Chestakova, Ronald J. Overmars, Stefan van Nieuwkoop, Richard Molenkamp, Annemiek van der Eijk, Corine GeurtsvanKessel, Harry Vennema, Adam Meijer, Andrew Rambaut, Jaap van Dissel, Reina S. Sikkema, Aura Timen, Marion Koopmans, on behalf of the Dutch-Covid-19 response team

**Author notes:** These authors contributed equally to this manuscript (Aura Timen, Marion Koopmans).

## Abstract

SARS-CoV-2 is a novel coronavirus that has rapidly spread across the globe. In the Netherlands, the first case of SARS-CoV-2 has been notified on the 27^th^ of February. Here, we describe the first three weeks of the SARS-CoV-2 outbreak in the Netherlands, which started with several different introductory events from Italy, Austria, Germany and France followed by local amplification in, and later also, outside the South of the Netherlands. The timely generation of whole genome sequences combined with epidemiological investigations facilitated early decision making in an attempt to control local transmission of SARS-CoV-2 in the Netherlands.

## Introduction

Late December 2019, a cluster of cases of pneumonia of unknown aetiology were reported in connection with a seafood and live animal market in Wuhan, China^1^. The virus responsible was tentatively named SARS-CoV-2^2^ and it rapidly spread throughout the world. By April 16^th^ the virus had spread to 185 different countries, infected over 2,000,000 people and resulted in over 130,000 deaths^3^.

Whole genome sequencing (WGS) is a powerful tool to understand transmission dynamics of (viral) outbreaks. Recently, several outbreaks of zoonotic viral pathogens have been reconstructed using genomic information^4–7^. Equally, WGS can be used to inform outbreak control decisions. Evidence of this was seen during the 2014-2016 West African Ebola outbreak when real-time WGS was used to help public health decision making, a strategy for which the term “precision public health pathogen genomics” has been dubbed^8,9^. In this definition, the use of WGS can provide a more detailed picture of the sources of infection and transmission chains. Immediate sharing and combining data during outbreaks is now recommended as an integral part of outbreak response^10–12^. Feasibility of real-time WGS relies on the access to sequence platforms that provide reliable sequences, access to metadata for interpretation, and data analysis at high speed and low cost. Therefore, whole genome sequencing for outbreak support is an active area of research. Nanopore sequencing has been employed in recent outbreaks of Usutu, Ebola, Zika and Yellow fever virus owing to the ease of use and relatively low start-up cost^4–7^. The robustness of this method has recently been validated using Usutu virus^13,14^.

In the Netherlands, the first COVID-19 case was confirmed on the 27^nd^ of February and as of April 16^th^, there were 29,214 SARS-CoV-2 positive patients and 3,351 patient deaths attributable to COVID-19 in the Netherlands. The first complete SARS-CoV-2 genome sequences of the first two patients in the Netherlands were generated, analyzed and shared on the 29^th^ of February. By March 15^th^, 189 full genome sequences were generated and released on GISAID. Here, we demonstrate how this data was generated and how SARS-CoV-2 WGS, in combination with epidemiological data, was used to inform public health decision making in the Netherlands.

## Methods

### COVID-19 response

This study was done in liaison with the national outbreak response team (ORT). The ORT develops guidance on case finding and containment, based on WHO and ECDC recommendations and expert advice, as defined by the crisis and emergency response structure^15,16^. The sequencing effort was embedded in the stepwise response to the outbreak (**Figure 1**), which evolved from the initial testing of symptomatic travelers according to the WHO and ECDC case definitions including the testing of symptomatic contacts (phase 1), followed by inclusion of routine testing of patients hospitalized with severe respiratory infections (phase 2), to inclusion of health care workers with a low threshold case definition and testing to define the extent of suspected clusters (phase 3). Depending on the phase and clinical severity, initial contact with patients was established through public health physicians or nurses from the municipal health service (for travel related cases, contacts of [hospitalized] cases, and patients belonging to risk groups). Ethical approval was not required for this study as only anonymous aggregated data were used, and no medical interventions were made on human subjects

**Figure 1:**
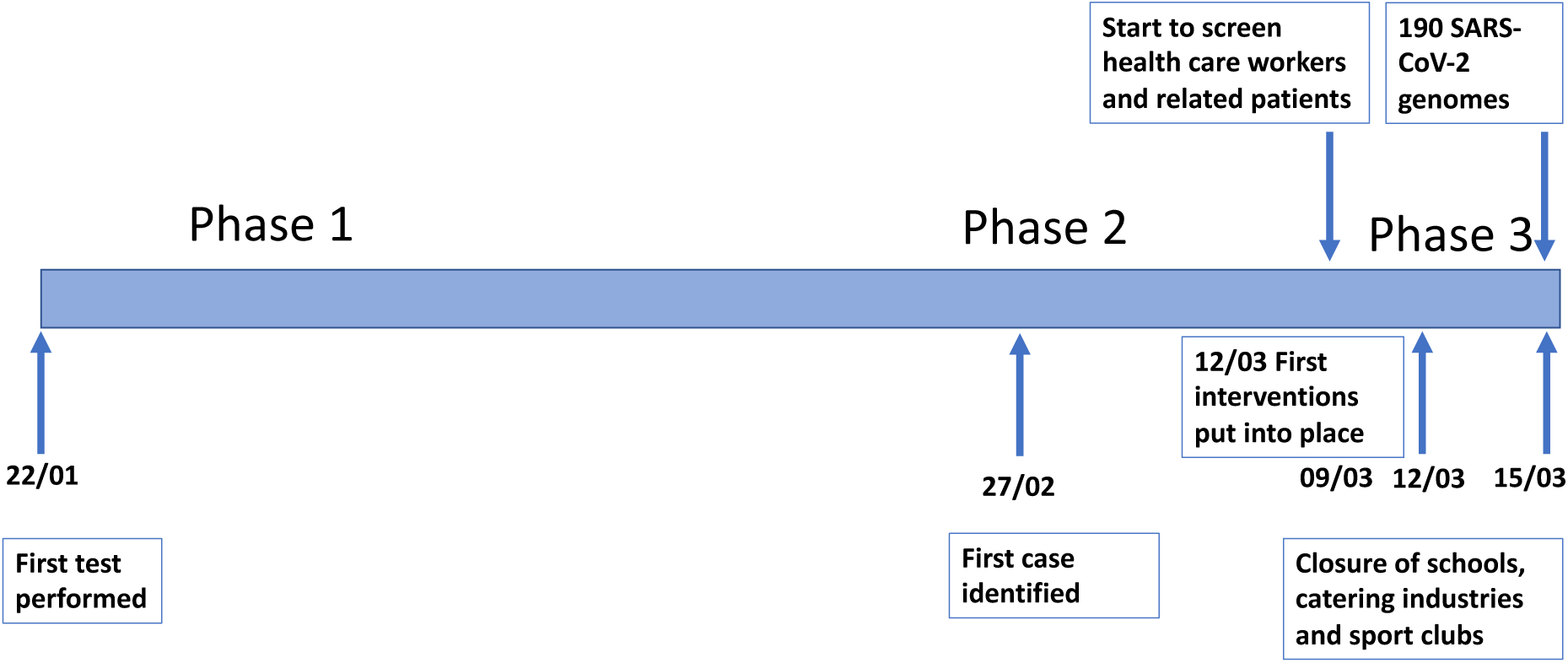
Timeline of the different phases in the response to the SARS-CoV-2 outbreak in the Netherlands.

### Contact tracing

On January 29th COVID-19 was classified as a notifiable disease in group A in the Netherlands, with physicians and laboratories having to report any suspected and confirmed case to the Dutch public health services (PHS) by phone. Upon notification, the PHS initiates source identification and contact tracing, and performs risk assessments. In the early outbreak phase (containment), the PHS traced and informed all high and low risk contacts of cases with the aim to stop further transmission. For each case, epidemiological information such as demographic information, symptoms, date of onset of symptoms, travel history, contact information, suspected source, underlying disease, and occupation were registered. Due to the magnitude of the COVID-19 outbreak this quickly became impracticable in severely affected regions, and the strategy shifted to registering data on confirmed cases only and informing their high-risk contacts (phase 2) with continued active case finding in less affected regions. The PHS informed the national public health authority of the Netherlands (RIVM) about all laboratory confirmed cases. There, a national case registry was kept in which a contact matrix was kept for the first 250 cases.

### SARS-CoV-2 diagnostics

Clinical specimens were collected and phocine distemper virus (PDV) was added as internal NA extraction control to the supernatant. Clinical specimens included oropharyngeal and nasopharyngeal swabs, bronchoalveolar lavage and sputum. Total NA was extracted from the supernatant using Roche MagNA Pure systems. The NA was screened for the presence of SARS-CoV-2 using real-time single-plex RT-PCRs for PDV, for the SARS-CoV-2 RdRp gene and for the SARS-CoV-2 E gene as described by Corman et al.^17^.

### SARS-CoV-2 whole genome sequencing

A SARS-CoV-2 specific multiplex PCR for Nanopore sequencing was performed, similar to amplicon-based approaches as previously described^18^. In short, primers for 86 overlapping amplicons spanning the entire genome were designed using primal^19^. The amplicon length was set to 500bp with 75bp overlap between the different amplicons. The used concentrations and primer sequences are displayed in **Supplementary table 1**. The libraries were generated using the native barcode kits from Nanopore (EXP-NBD104 and EXP-NBD114 and SQK-LSK109) and sequenced on a R9.4 flow cell multiplexing up to 24 samples per sequence run.

In the first phase all samples were selected for sequencing, reflecting travel associated cases and their contacts. In the second phase priority was given to patients identified through enhanced case finding by testing of hospitalized patients with severe acute respiratory disease (SARI) and continued sequencing of new incursions. In the third phase, the epidemic started to expand exponentially, and sequencing was done to continue to monitor evolution of the outbreak. In line with the national testing policy, a significant proportion of new cases sequenced were healthcare workers.

### Sequence data analysis

The resulting raw sequence data was demultiplexed using qcat (https://github.com/nanoporetech/qcat) or Porechop (https://github.com/rrwick/Porechop). Primers were trimmed using cutadapt^20^ after which a reference based alignment was performed using minimap2^21^ to GISAID sequence EPI_ISL_412973. The run was monitored using RAMPART (https://artic-network.github.io/rampart/) and the analysis process was automated using snakemake^22^ which was used to perform near to real-time analysis with new data every 10 minutes. The consensus genome was extracted and positions with a coverage <30 were replaced with an “N” with a custom script using biopython and pysam. Mutations in the genome were confirmed by manually checking the alignment and homopolymeric regions were manually checked and resolved consulting reference genomes.

### Phylogenetic analysis

All available full-length SARS-CoV-2 genomes were retrieved from GISAID (**supplementary table 2**) on the 22^nd^ of March 2020 and aligned with the Dutch SARS-CoV-2 sequences from this study using MUSCLE. Sequences with >10% “Ns” were excluded. The alignment was manually checked for discrepancies after which IQ-TREE^23^ was used to perform a maximum likelihood phylogenetic analysis under the GTR+F+I +G4 model as best predicted model using the ultrafast bootstrap option with 1,000 replicates. The phylogenetic trees were visualized using custom python and baltic scripts (https://github.com/evogytis/baltic).

### BEAST analysis

All available full-length SARS-CoV-2 genomes were retrieved from GISAID^24,25^ on the 18th of March 2020 and downsampled to include only representative sequences from epidemiologically linked cases. Sequences lacking date information were also removed from the dataset. Sequences were aligned with the newly obtained SARS-CoV-2 sequences in this study using MAFFT^26^. Bayesian phylogenetic trees were inferred using BEAST^27^. An exponential growth model and skygrid model were set to 100,000,000 states with sampling every 10,000 states. Log files were inspected in Tracer and Tree annotator was used to remove the burn-in from the tree file and to construct the mcc tree.

## Results

After the release of the first SARS-CoV-2 genomic sequence, a diagnostic assay was quickly set-up and validated^17^. Diagnostics were initially performed on suspected cases with a recent travel history to China, but later also on suspected cases with a recent travel history to parts of Italy, Germany, Austria or France. In addition, an amplicon-based full genome sequencing protocol was developed based on the release of the first genomic sequence.

### Phase 1: Initial testing of travelers according to the WHO and ECDC case definitions

Starting from the 30th of January symptomatic travelers from countries where SARS-CoV-2 was known to circulate were routinely tested. This resulted in the identification of the first SARS-CoV-2 infection in the Netherlands on February 27^th^ and an additional case one day later. The genomes of these first two positive samples were generated and analyzed by the February 29^th^. These two viruses clustered differently in the phylogenetic tree, confirming separate introductions (**Figure 2A**).

**Figure 2:**
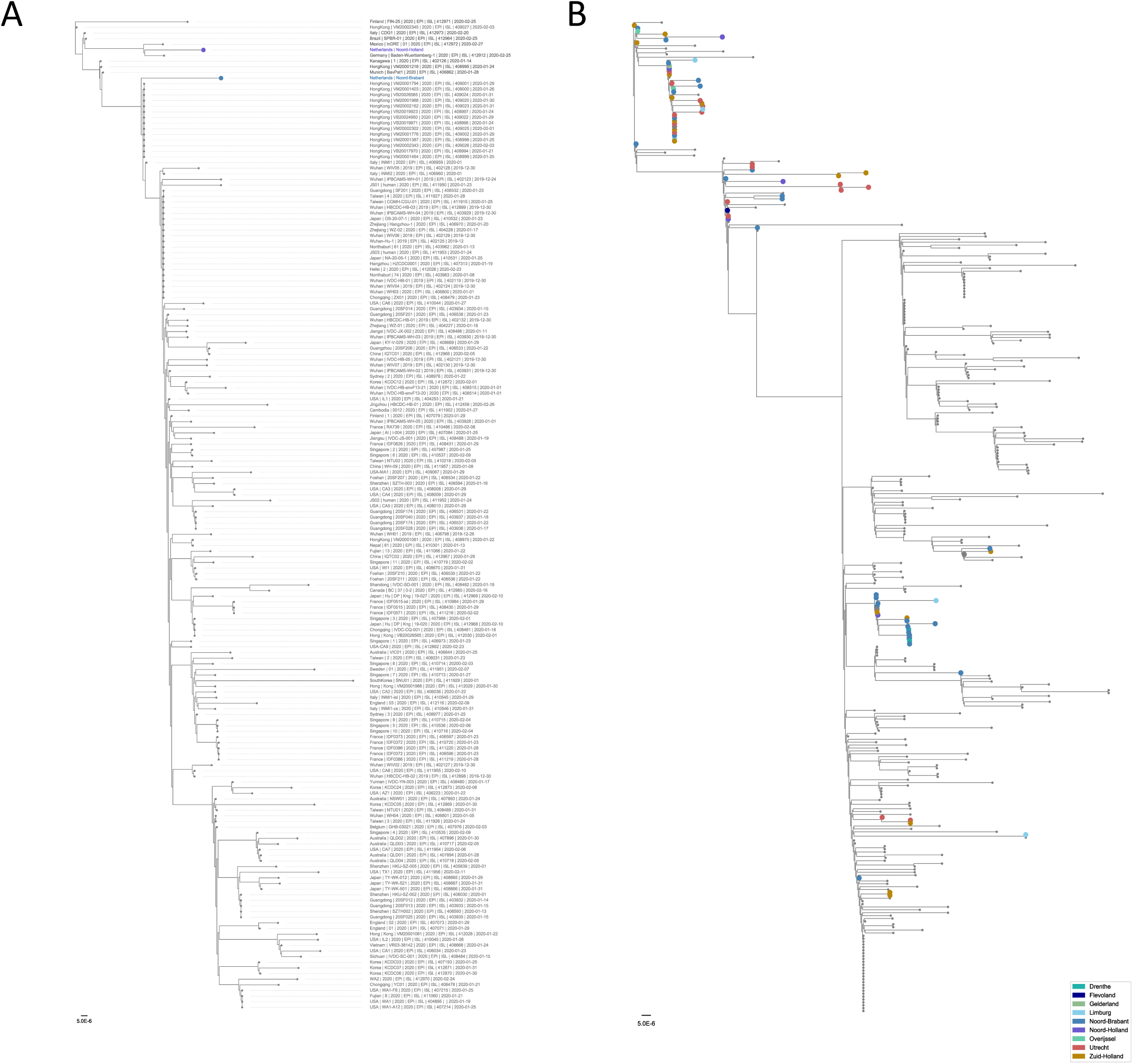
Phylogenetic analysis of the first two Dutch SARS-CoV-2. All sequences that were publicly available on the 29^th^ of February (panel A) or on the 9^th^ of March (panel B) are included in the analysis. Sequences are colored based on province of detection. Scale bar represents the amount of nucleotide substitutions per site. Red indicates the Dutch isolates and blue represent SARS-CoV-2 sequences from other countries with recent travel history to Italy.

During the following days the number of reported cases rapidly increased to 321 through screening of symptomatic travelers and symptomatic contacts. Also, more sequence data was generated: by March 9^th^ local clusters of epidemiologically related cases of SARS-CoV-2 started to appear in the province of Noord Brabant. The increase in cases is likely to have been triggered by multiple introductions of the virus following the spring holidays (from 13^th^ of February to 23^rd^ of February) with travel to ski resorts in Northern Italy (**Figure 2B**). The virus also started to appear in other parts of the country (**Figure 3A**) through which on March 9^th^ the first intervention was put into place: the prime minister advised people to stop shaking hands. One day later, on the 10^th^ of March, events attended by more than 1,000 visitors were banned in the province North Brabant.

**Figure 3:**
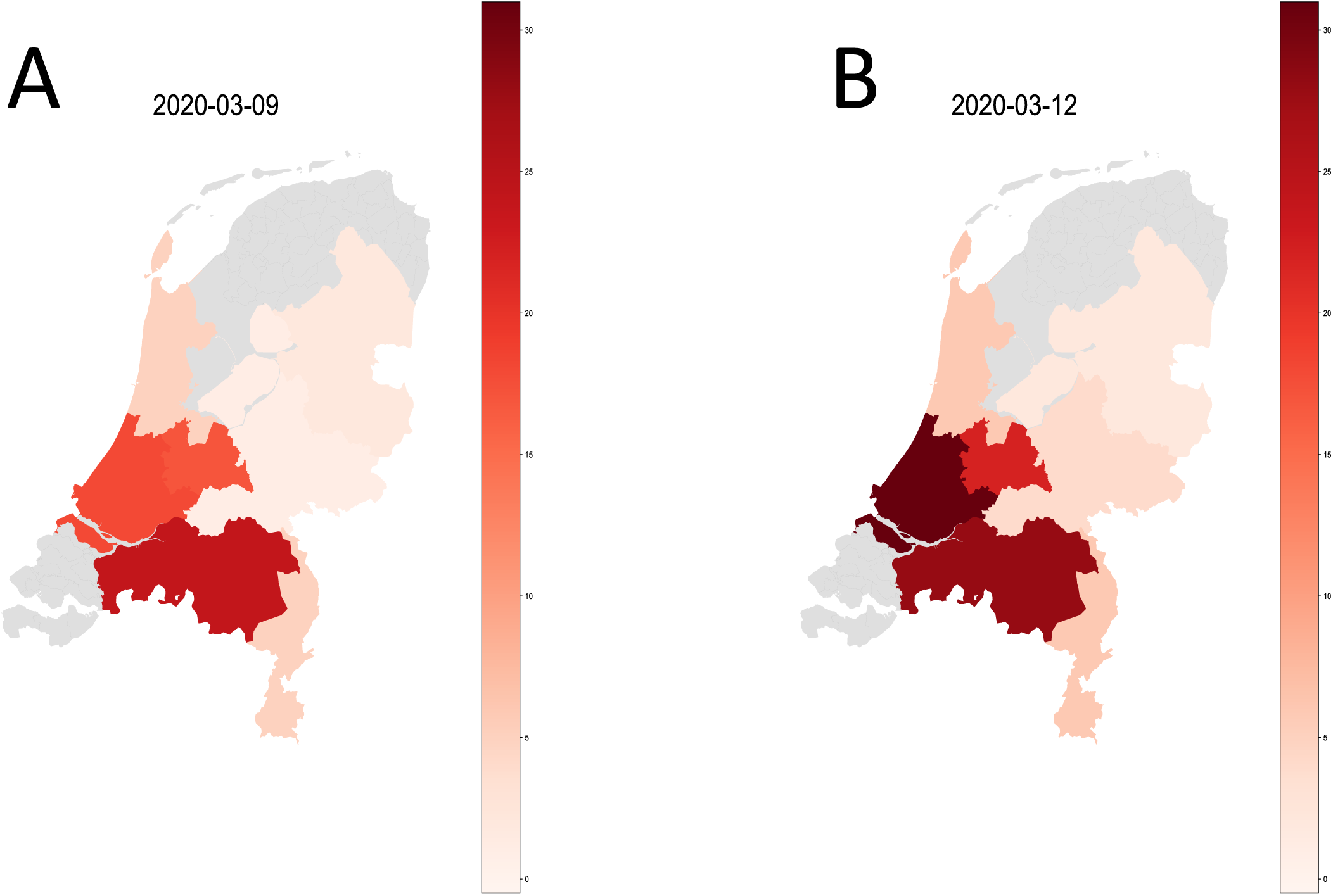
Distribution of SARS-CoV-2 sequences from the Netherlands on the 9^th^ of March and the 12^th^ of March and the 15^th^ of March. The source of the shapefile of the map is derived from gadm.org.

### Phase 2: Inclusion of patients hospitalized with severe respiratory infections

The advice to test hospitalized patients with serious respiratory infections was issued on February 24^rd^ and subsequent attempts to identify possible local transmission chains triggered testing for SARS-CoV-2 on a large scale in hospitals. All positive samples were sequenced and during the following days, whole genome sequencing and subsequent phylogenetic analysis revealed the emergence of multiple viral strains that clustered differently in the phylogenetic tree as well as substantial local amplification of viruses from patients without any travel history, also outside Noord Brabant (**Figure 3B**). This information, combined with the increase in the total number of infections in the Netherlands, led to the decision to implement stricter measures for the whole country to prevent further spread of SARS-CoV-2 on the 12^th^ of March. All events with over 100 people attending were cancelled, people were requested to work from home as much as possible and people with symptoms like fever or cough had to stay at home. On the 15^th^ of March this was followed by the closure of schools, catering industries and sport clubs.

### Phase 3: Systematic sequencing during exponential growth phase

In the third phase sequencing of new cases with emphasis on HCW and severe hospitalized cases was continued. By March 15^nd^, 190 SARS-CoV-2 viruses from the Netherlands were sequenced, at that moment representing 27,1% of the total number of full genome sequences produced worldwide. The sequences detected in the Netherlands were diverse and revealed the presence of multiple co-circulating sequence types, found in several different clusters in the phylogenetic tree (**Figure 4 and Supplementary Figure 1**). The diversity was also observed in cases with similar travel histories, reflecting that sequence diversity already was present in the originating county, primarily Italy (**Figure 4**). In addition to travel associated cases, an increasing number of local cases was detected through SARI surveillance; this was not limited to the province Noord Brabant but SARS-CoV-2 was also increasing in the provinces Zuid Holland, Noord Holland and Utrecht, confirming substantial under ascertainment of the epidemic. We also observed the occurrence of clusters, some of which triggered in depth investigations to be described elsewhere (Sikkema et al., in preparation). The increase in COVID-19 patients as well as increasing affected geographic areas and occurrence of local clusters provided further support for the increased movement restrictions.

**Figure 4:**
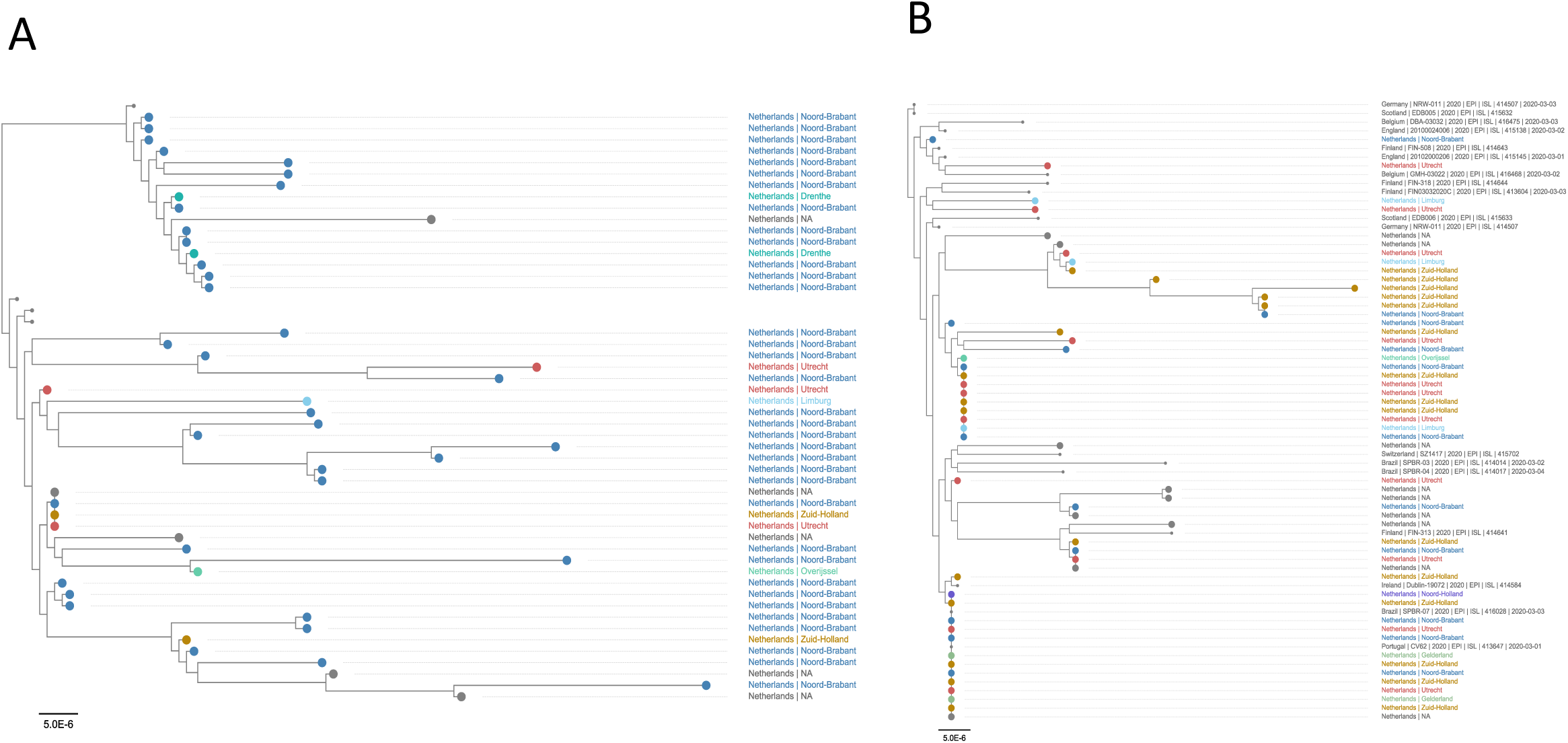
Phylogenetic analysis of SARS-CoV-2 emergence in the Netherlands. Panel A and B represent a zoom-in of two clusters circulating in the Netherlands. In Panel A only names from sequences from the Netherlands are indicated while in panel B all names are indicated. Sequences are colored based on province of detection. Scale bar represent the number of substitutions per site.

### Time-resolved analysis

BEAST analysis revealed that the most recent ancestor of the viruses circulating in the Netherlands dates back to the end of January and the beginning of February (**Figure 5**). This is in line with the amplification that occurred in the region (notably Italy and Switzerland) from which most of the epidemic in The Netherlands was seeded. Most incursions likely occurred during spring break, which is a popular winter sports vacation area. Retrospective testing showed the presence of the virus in a sample collected on the 24^th^ of February in a patient with known travel history to Italy.

**Figure 5:**
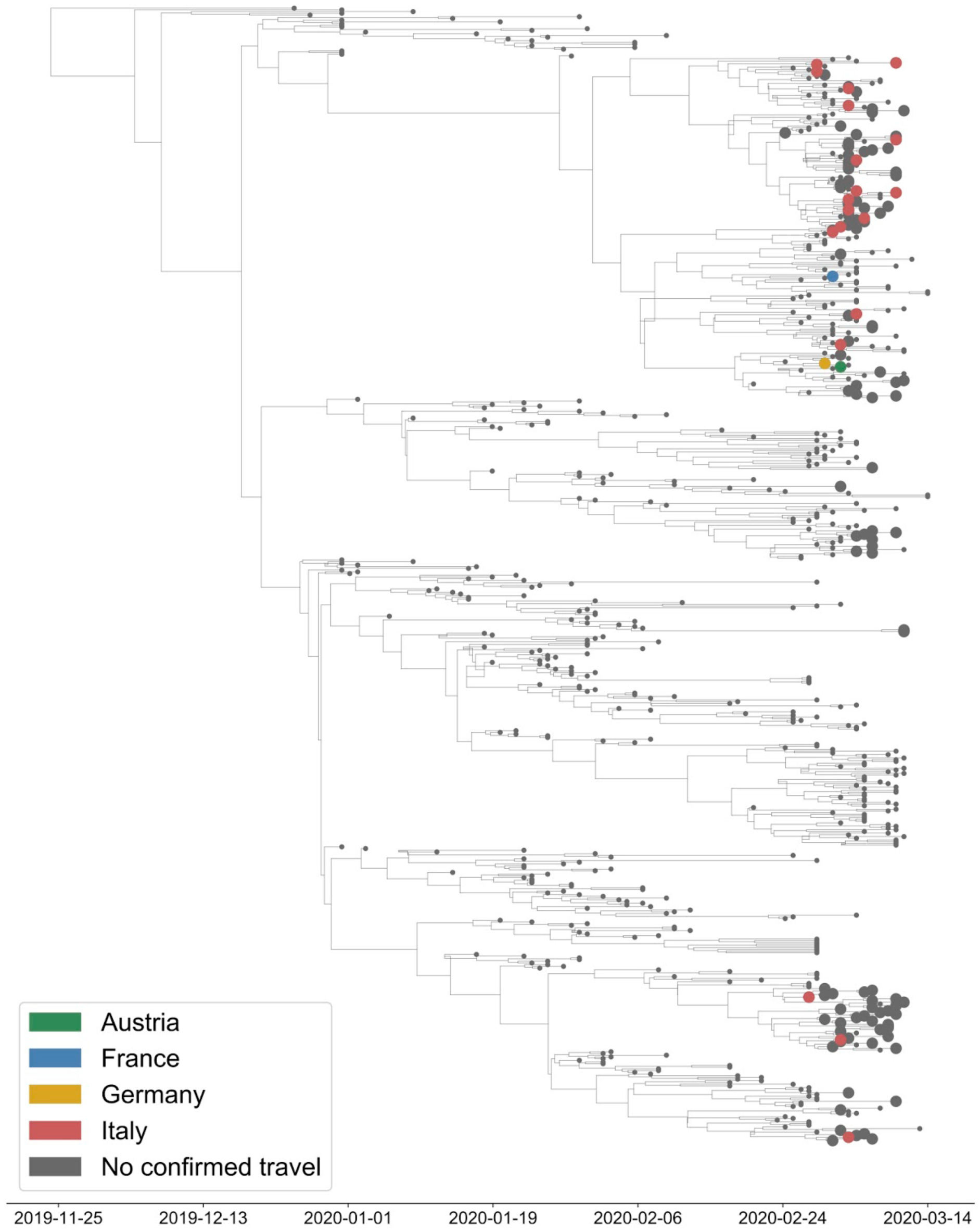
BEAST analysis with travel history. Time resolved visualization of the emergence of SARS-CoV-2 in the Netherlands. Sequences from the Netherlands are depicted in big circles. Green indicates recent travel history to Austria, blue to France yellow to Germany, and red to Italy.

## Discussion

We show that whole genome sequencing in combination with epidemiological data strengthened the evidence base for public health decision making in the Netherlands. WGS enabled a more precise understanding of the transmission patterns in various initial phases of the outbreaks. As such, we were able to understand the genetic diversity of the multiple introduction events in phase 1, the extent of local and regional clusters in phase 2 and the transmission patterns within the HCW groups in phase 3 (among which the absence or occurrence of very limited nosocomial transmission). This information complemented the data obtained from more traditional methods such as contact investigation. To yield optimal value for public health measures, WGS was performed on the basis of questions raised by the multidisciplinary national response team. This timely generation of whole genome sequences facilitated early decision making in an attempt to control local transmission of SARS-CoV-2 in the Netherlands.

At the moment, sequences from the Netherlands make up a substantial part of the total collection of SARS-CoV-2 genomes. Although implementation of WGS in the Dutch disease prevention and control has shown its added value, there is still only a limited amount of genomic information available from certain parts of the world, including Italy, and therefore, conclusions need to be drawn with caution given the inevitable sampling bias of the dataset. Also, from Iran, another major country where the virus presumably was spreading exponentially in the week before the take-off of the epidemic in The Netherlands, only genomic information about exported cases is known. Without a representative and sizable selection of reference sequences, reliable phylogenetic analysis is difficult for the time being, although the geographic signature in the dataset is becoming increasingly clear. Clustering, and conclusions on the origin of viruses may change significantly when virus sequences of other geographical regions are added to the analysis. Moreover, global monitoring of the genetic diversity of the virus is essential to reliably model and predict the spread of the virus.

Nonetheless, there are a few observations that can be made: since its emergence, the global spread of SARS-CoV-2 led to diversification into lineages that reflect ongoing chains of transmission in specific geographic regions globally, in Europe, and -during the 2nd and 3d phase-in the Netherlands. This diversification provided the basis for the use of WGS to investigate possible transmission chains locally, for instance in healthcare settings, where it can be used to inform infection control and prevention when combined with background data on contact histories among others. Moreover, the continued effort will lay the foundation for the enhanced surveillance that will be paramount during the next phase of the pandemic, when confinement measures will gradually be lifted. Given the widespread circulation, most likely scenario is that SARS-CoV-2 will (sporadically) re-emerge, and discrimination between novel introductions versus prolonged local circulation is important to inform appropriate public health decision. In addition, due to genomic mutations, the phenotype and the transmission dynamics of the virus might change during time. Therefore, close monitoring of the behavior of the virus in combination with genetic information is essential as well.

We have used an amplicon-based sequencing approach to monitor the emergence of SARS-CoV-2 in the Netherlands. A critical step in using amplicon-based sequencing is that close, reliable reference sequences need to be available. The primers are designed based on our current knowledge about SARS-CoV-2 diversity and therefore need regular updating. In the future, this may be overcome using metagenomic sequencing. However, at the moment conventional metagenomic sequencing (Illumina) takes too long for near to real-time sequencing and Nanopore based metagenomic sequencing is not sensitive enough to allow recovery of whole genome sequences in a similar fashion and similar costs as compared to amplicon based Nanopore sequencing.

We provide a first description of the incursion of SARS-CoV-2 into The Netherlands. The combination of real-time WGS with the data from the National Public Health response team has provided information that helped decide on next steps in the decision making. Sharing of metadata is needed within a country but also on a global level. We urge countries to share sequence information to combine our efforts in understanding the spread of SARS-CoV-2. GISAID^24,25^ made sharing of sequence information coupled to limited metadata possible in a manner that protects the intellectual property and acknowledges the data providers. However, in order to fully capitalize on the potential added value of WGS for public health decision making, systems for combined analysis of data are needed that are in agreement with general data protection rules. We previously developed a model for collaborative exploration of WGS and metadata in a protected sharing environment^28,29^ For truly global collaboration, such systems would need to be further developed and hosted under the auspices of WHO.

## Acknowledgements

We gratefully acknowledge the originating laboratories, where specimens were first obtained, and the submitting laboratories, where sequence data were generated and submitted to the EpiFlu Database of the Global Initiative on Sharing All Influenza Data (GISAID), on which this research is based. All contributors of data may be contacted directly via the GISAID website (http://platform.gisaid.org). The accession numbers of all genetic sequences used in this study are provided in **supplementary table 2** and are accessible from the website of GISAID (http://platform.gisaid.org).

## Dutch-Covid-19 response team

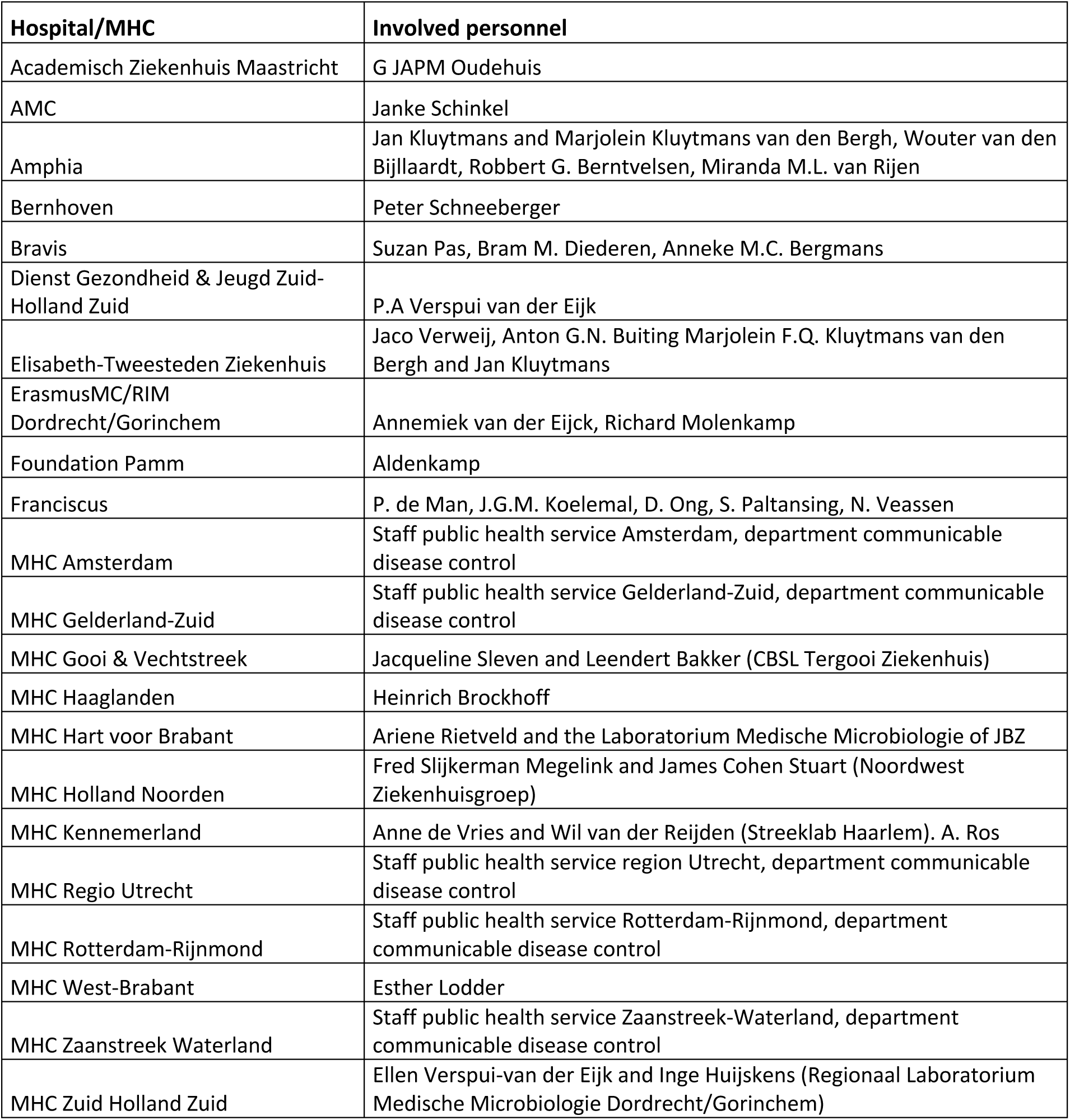
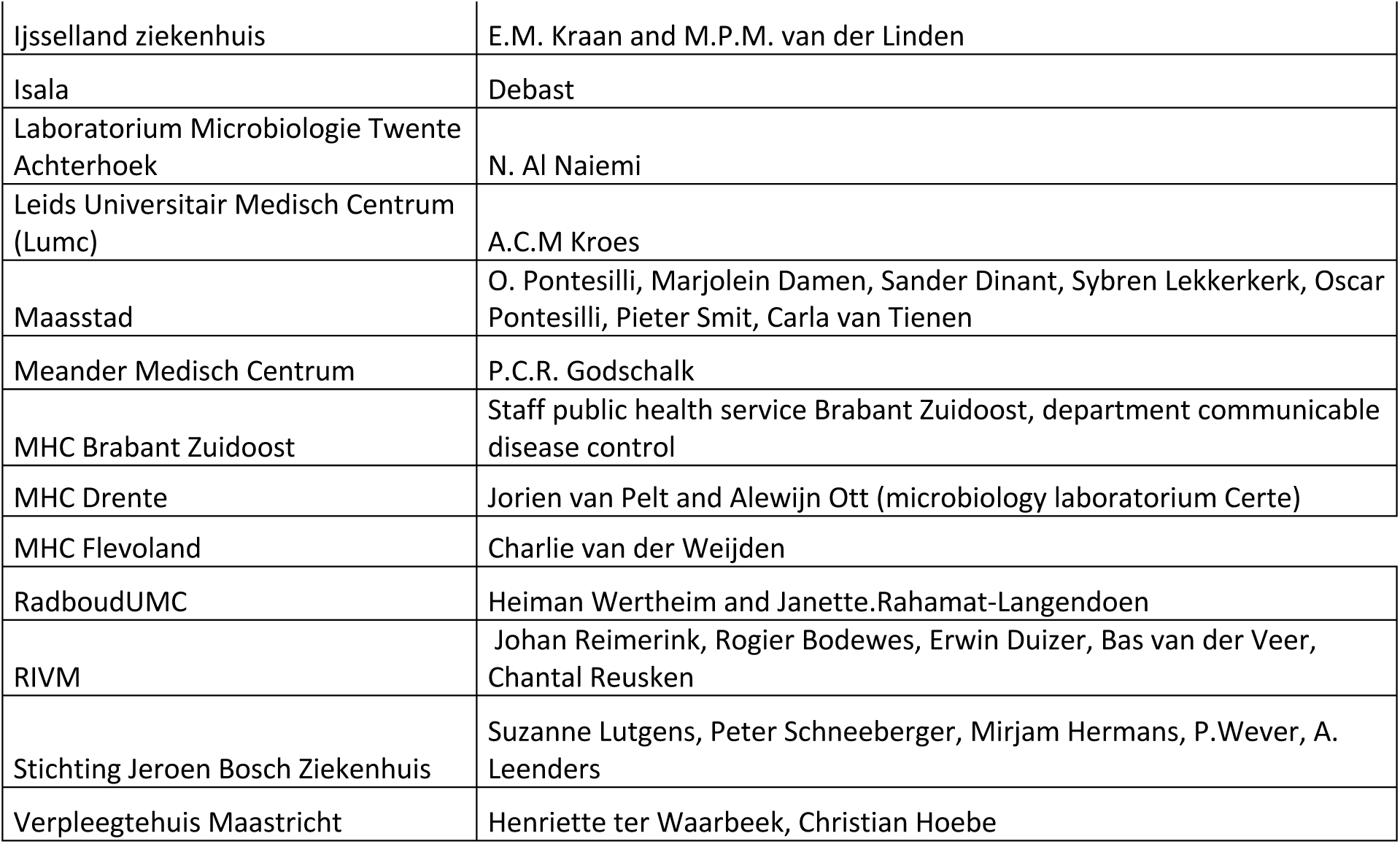

**Supplementary figure 1:** Full maximum likelihood tree. Sequences are colored based on province of detection. Scale bar represent the number of substitutions per site.

**Supplementary table 1:** Primer sequences and volumes as used in this study. Primers were dissolved to a concentration of 100 μM.

**Supplementary table 2:** GISAID acknowledgements table

