## Supplementary figures and images for "Rapid SARS-CoV-2 whole genome sequencing for informed public health decision making in the Netherlands"

### Supplementary figure 1

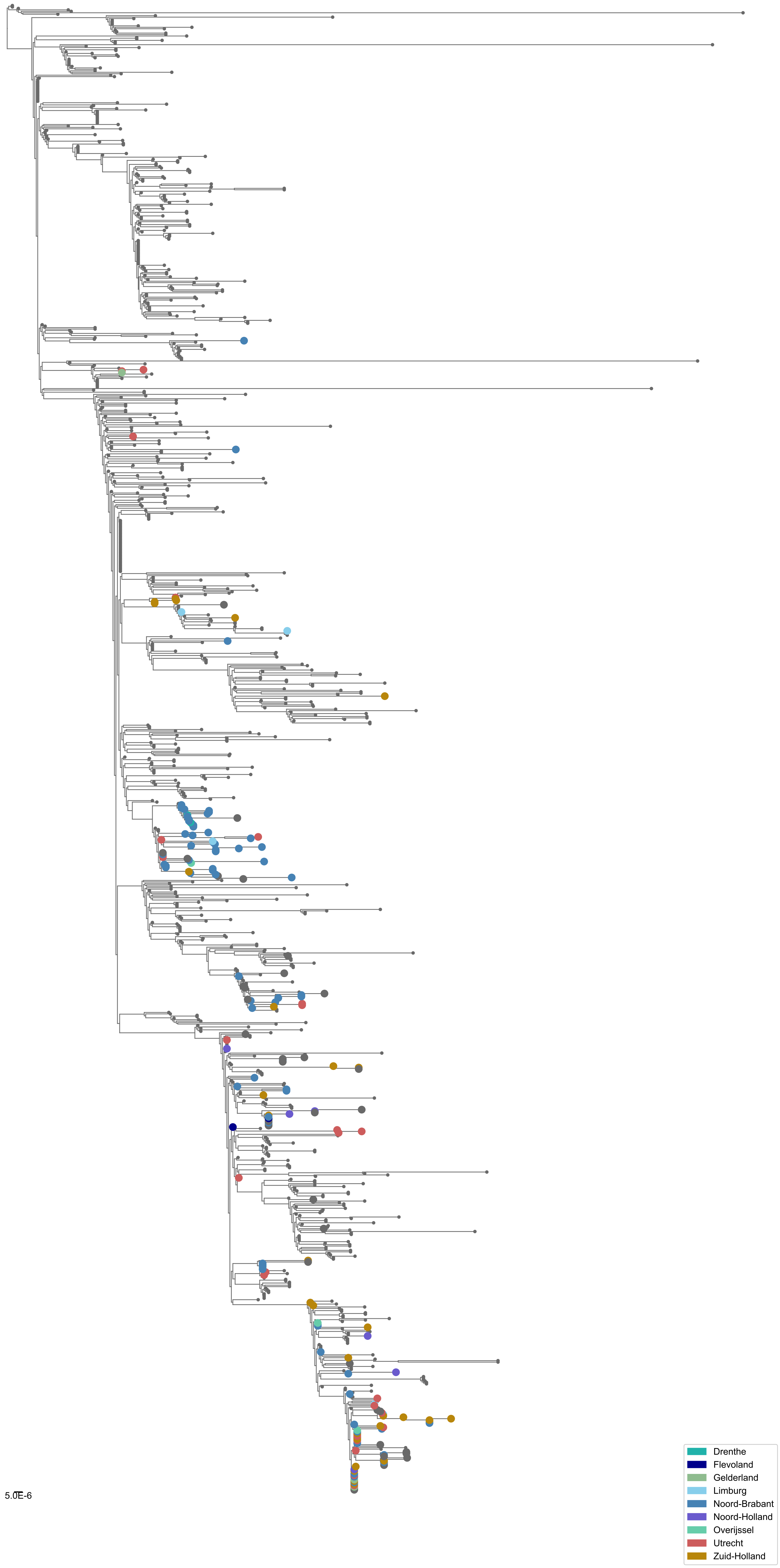
