## Supplementary table 1 for "Rapid SARS-CoV-2 whole genome sequencing for informed public health decision making in the Netherlands"

**Supplementary table 1:** Primer sequences and volumes as used in this study. Primers were dissolved to a concentration of 100 μM.

| **Mix 1** |  |  |  |
| --- | --- | --- | --- |
| **Name** | **Sequence** | **Orientation:** | **Volume:** |
| SARS-CoV-2_1_LEFT | ACCAACCAACTTTCGATCTCTTGT | FWD | 10μl |
| SARS-CoV-2_1_RIGHT | CGAGCATCCGAACGTTTGATGA | REV | 10μl |
| SARS-CoV-2_3_LEFT | ACGAGCTTGGCACTGATCCTTA | FWD | 10μl |
| SARS-CoV-2_3_RIGHT | GGTTGCATTCATTTGGTGACGC | REV | 10μl |
| SARS-CoV-2_5_LEFT | TGTCCAGCATGTCACAATTCAGA | FWD | 10μl |
| SARS-CoV-2_5_RIGHT | TACAACACGAGCAGCCTCTGAT | REV | 10μl |
| SARS-CoV-2_7_LEFT | TGGCACTGTTTATGAAAAACTCAAACC | FWD | 10μl |
| SARS-CoV-2_7_RIGHT | TTTCGAGCAACATAAGCCCGTT | REV | 10μl |
| SARS-CoV-2_9_LEFT | TCACTTTTGAACTTGATGAAAGGATTGA | FWD | 10μl |
| SARS-CoV-2_9_RIGHT | GATTGTCCTCACTGCCGTCTTG | REV | 10μl |
| SARS-CoV-2_11_LEFT | AAACATGGAGGAGGTGTTGCAG | FWD | 20μl |
| SARS-CoV-2_11_RIGHT | TCTTGTTTTCTCTGTTCAACTGAAGGT | REV | 20μl |
| SARS-CoV-2_13_LEFT | GCTCCATATATAGTGGGTGATGTTGT | FWD | 10μl |
| SARS-CoV-2_13_RIGHT | AGCCATGTGTTACATAGCCAAGTG | REV | 10μl |
| SARS-CoV-2_15_LEFT | CTATTCTGGACAATCTACACAACTAGGT | FWD | 10μl |
| SARS-CoV-2_15_RIGHT | AGCATCTTGTAGAGCAGGTGGA | REV | 10μl |
| SARS-CoV-2_17_LEFT | GCTGTTATGTACATGGGCACACT | FWD | 10μl |
| SARS-CoV-2_17_RIGHT | GCTTGCGTTTGGATATGGTTGG | REV | 10μl |
| SARS-CoV-2_19_LEFT | TGTTACATAAACCTATTGTTTGGCATGT | FWD | 10μl |
| SARS-CoV-2_19_RIGHT | TCCCAAGGGACACTATTAACAGCA | REV | 10μl |
| SARS-CoV-2_21_LEFT | AGCAAAGAATACTGTTAAGAGTGTCGG | FWD | 20μl |
| SARS-CoV-2_21_RIGHT | TCGGGGCCATTTGTACAAGATT | REV | 20μl |
| SARS-CoV-2_23_LEFT | ACTATTGTTAATGGTGTTAGAAGGTCCT | FWD | 10μl |
| SARS-CoV-2_23_RIGHT | GCAACTTCCGCACTATCACCAA | REV | 10μl |
| SARS-CoV-2_25_LEFT | TGTCTTAAATTGTCACATCAATCTGACAT | FWD | 10μl |
| SARS-CoV-2_25_RIGHT | TCACGAGTGACACCACCATCAA | REV | 10μl |
| SARS-CoV-2_27_LEFT | CTGGTTTGCCTGGCACGATATT | FWD | 10μl |
| SARS-CoV-2_27_RIGHT | TCTACACCACAGAAAACTCCTGGT | REV | 10μl |
| SARS-CoV-2_29_LEFT | ACAGTCATGTAGTTGCCTTTAATACTTTAC | FWD | 10μl |
| SARS-CoV-2_29_RIGHT | GAGCCTTTGCGAGATGACAACA | REV | 10μl |
| SARS-CoV-2_31_LEFT | AACGGTCTTTGGCTTGATGACG | FWD | 10μl |
| SARS-CoV-2_31_RIGHT | ACCTTCTAAGTCTGTGCCAGCA | REV | 10μl |
| SARS-CoV-2_33_LEFT | TGTGGCTATGAAGTACAATTATGAACCT | FWD | 10μl |
| SARS-CoV-2_33_RIGHT | AGCTACAGTGGCAAGAGAAGGT | REV | 10μl |
| SARS-CoV-2_35_LEFT | TGGTGCTAGGAGAGTGTGGACA | FWD | 10μl |
| SARS-CoV-2_35_RIGHT | GGCTACTTTGATACAAGGTTTGCC | REV | 10μl |
| SARS-CoV-2_37_LEFT | CTTTCCATGCAGGGTGCTGTAG | FWD | 10μl |
| SARS-CoV-2_37_RIGHT | GTTTGGCTGCTGTTGTAAGAGGT | REV | 10μl |
| SARS-CoV-2_39_LEFT | ACAATTCACCTAATTTAGCATGGCCT | FWD | 10μl |
| SARS-CoV-2_39_RIGHT | TTGGTTGTCCCCCACTAGCTAG | REV | 10μl |
| SARS-CoV-2_41_LEFT | AGTATGTACAAATACCTACAACTTGTGCT | FWD | 10μl |
| SARS-CoV-2_41_RIGHT | AGCATAGACGAGGTCTGCCATT | REV | 10μl |
| SARS-CoV-2_43_LEFT | TGGTATTGTTGGTGTACTGACATTAGA | FWD | 10μl |
| SARS-CoV-2_43_RIGHT | AGGGTCAGCAGCATACACAAGT | REV | 10μl |
| SARS-CoV-2_45_LEFT | CACTTCTTCTTTGCTCAGGATGGT | FWD | 10μl |
| SARS-CoV-2_45_RIGHT | AACATGTTGTGCCAACCACCAT | REV | 10μl |
| SARS-CoV-2_47_LEFT | TGTAGCTTGTCACACCGTTTCT | FWD | 10μl |
| SARS-CoV-2_47_RIGHT | AGTAAGGTCAGTCTCAGTCCAACA | REV | 10μl |
| SARS-CoV-2_49_LEFT | AGGAGTATGCTGATGTCTTTCATTTGT | FWD | 10μl |
| SARS-CoV-2_49_RIGHT | CACCAGCATTTGTCCAGTCACA | REV | 10μl |
| SARS-CoV-2_51_LEFT | TCTTTCATGGGAAGTTGGTAAACCT | FWD | 20μl |
| SARS-CoV-2_51_RIGHT | TCACATAGTGCATCAACAGCGG | REV | 20μl |
| SARS-CoV-2_53_LEFT | TGTCAATGCCAGATTAYGTGCT | FWD | 10μl |
| SARS-CoV-2_53_RIGHT | CAAGAGTGAGCTGTTTCAGTGGT | REV | 10μl |
| SARS-CoV-2_55_LEFT | TGTTGACACTAAATTCAAAACTGAAGGT | FWD | 20μl |
| SARS-CoV-2_55_RIGHT | TGTCAACTCAAAGCCATGTGCC | REV | 20μl |
| SARS-CoV-2_57_LEFT | CGTTTATGATTGATGTTCAACAATGGGG | FWD | 10μl |
| SARS-CoV-2_57_RIGHT | ACAACCAGGCAAGTTAAGGTTAGA | REV | 10μl |
| SARS-CoV-2_59_LEFT | AGTCTCATGGAAAACAAGTAGTGTCA | FWD | 10μl |
| SARS-CoV-2_59_RIGHT | ATTAGCAGCAATGTCCACACCC | REV | 10μl |
| SARS-CoV-2_61_LEFT | AGAAATGCCCGTAATGGTGTTCT | FWD | 10μl |
| SARS-CoV-2_61_RIGHT | TGAACCTGTTTGCGCATCTGTT | REV | 10μl |
| SARS-CoV-2_63_LEFT | TGGCCATGTAGAAACATTTTACCCA | FWD | 10μl |
| SARS-CoV-2_63_RIGHT | ATAGCCACGGAACCTCCAAGAG | REV | 10μl |
| SARS-CoV-2_65_LEFT | GGCAAACCACGCGAACAAATAR | FWD | 10μl |
| SARS-CoV-2_65_RIGHT | ACCTCTTAGTACCATTGGTCCCA | REV | 10μl |
| SARS-CoV-2_67_LEFT | CAATTTTGTAATGATCCATTTTTGGGTGT | FWD | 10μl |
| SARS-CoV-2_67_RIGHT | GGTCAAGTGCACAGTCTACAGC | REV | 10μl |
| SARS-CoV-2_69_LEFT | ACTGTGTTGCTGATTATTCTGTCCT | FWD | 20μl |
| SARS-CoV-2_69_RIGHT | TAGGTCCACAAACAGTTGCTGG | REV | 20μl |
| SARS-CoV-2_71_LEFT | ACTTCTAACCAGGTTGCTGTTCTT | FWD | 10μl |
| SARS-CoV-2_71_RIGHT | CAGCTATTCCAGTTAAAGCACGGT | REV | 10μl |
| SARS-CoV-2_73_LEFT | CTTGCAGATGCTGGCTTCATCA | FWD | 10μl |
| SARS-CoV-2_73_RIGHT | TGCACTTCAGCCTCAACTTTGT | REV | 10μl |
| SARS-CoV-2_75_LEFT | CTTCCCTCAGTCAGCACCTCAT | FWD | 10μl |
| SARS-CoV-2_75_RIGHT | CAAGCCAGCTATAAAACCTAGCCA | REV | 10μl |
| SARS-CoV-2_77_LEFT | TGGAACTGTAACTTTGAAGCAAGGT | FWD | 10μl |
| SARS-CoV-2_77_RIGHT | GACTTGTTGTGCCATCACCTGA | REV | 10μl |
| SARS-CoV-2_79_LEFT | GGTGTTGAACATGTTACCTTCTTCAT | FWD | 10μl |
| SARS-CoV-2_79_RIGHT | GTACCGTTGGAATCTGCCATGG | REV | 10μl |
| SARS-CoV-2_81_LEFT | CTTGTTTTGTGCTTGCTGCTGT | FWD | 10μl |
| SARS-CoV-2_81_RIGHT | ACTGCTACTGGAATGGTCTGTGT | REV | 10μl |
| SARS-CoV-2_83_LEFT | TGAAGAGCAACCAATGGAGATTGA | FWD | 10μl |
| SARS-CoV-2_83_RIGHT | TGTTCGTTTAGGCGTGACAAGT | REV | 10μl |
| SARS-CoV-2_85_LEFT | AGCACCTTTAATTGAATTGTGCGTG | FWD | 10μl |
| SARS-CoV-2_85_RIGHT | CGTCTGGTAGCTCTTCGGTAGT | REV | 10μl |
| SARS-CoV-2_87_LEFT | AAAAGATCACATTGGCACCCGC | FWD | 10μl |
| SARS-CoV-2_87_RIGHT | CGACATTCCGAAGAACGCTGAA | REV | 10μl |
| SARS-CoV-2_89_LEFT | AGGCTGATGAAACTCAAGCCTT | FWD | 20μl |
| SARS-CoV-2_89_RIGHT | AAAATCACATGGGGATAGCACTACT | REV | 20μl |

| **Mix 2** |  |  |  |
| --- | --- | --- | --- |
| **Name** | **Sequence** | **Orientation:** | **Volume:** |
| SARS-CoV-2_2_LEFT | TCGTACGTGGCTTTGGAGACTC | FWD | 10μl |
| SARS-CoV-2_2_RIGHT | ATGCACTCAAGAGGGTAGCCAT | REV | 10μl |
| SARS-CoV-2_4_LEFT | ACACCTTCAATGGGGAATGTCC | FWD | 10μl |
| SARS-CoV-2_4_RIGHT | AGGCACACTTGTTATGGCAACC | REV | 10μl |
| SARS-CoV-2_6_LEFT | TGTGAAAGGTTTGGATTATAAAGCATTCA | FWD | 10μl |
| SARS-CoV-2_6_RIGHT | ACAGGTGACAATTTGTCCACCG | REV | 10μl |
| SARS-CoV-2_8_LEFT | AGGGAGAAACACTTCCCACAGA | FWD | 10μl |
| SARS-CoV-2_8_RIGHT | AATCAATGCCCAGTGGTGTAAGT | REV | 10μl |
| SARS-CoV-2_10_LEFT | TGAGTATGGTACTGAAGATGATTACCAAG | FWD | 10μl |
| SARS-CoV-2_10_RIGHT | GCCGACAACATGAAGACAGTGT | REV | 10μl |
| SARS-CoV-2_12_LEFT | ACTGTTCGCACGAAYGTCTACT | FWD | 10μl |
| SARS-CoV-2_12_RIGHT | CCTGACCCGGGTAAGTGGTTAT | REV | 10μl |
| SARS-CoV-2_14_LEFT | GTTTCAACTATACAGCGTAAATATAAGGGT | FWD | 20μl |
| SARS-CoV-2_14_RIGHT | CGTGTGGAGGTTAATGTTGTCTACT | REV | 20μl |
| SARS-CoV-2_16_LEFT | AGGTACATGTCAGCATTAAATCACACT | FWD | 10μl |
| SARS-CoV-2_16_RIGHT | AGTTCATACTGAGCAGGTGGTG | REV | 10μl |
| SARS-CoV-2_18_LEFT | CAGTTACACAACAACCATAAAACCAGT | FWD | 10μl |
| SARS-CoV-2_18_RIGHT | GATTATCCATTCCCTGCGCGTC | REV | 10μl |
| SARS-CoV-2_20_LEFT | TACAGAAGAGGTTGGCCACACA | FWD | 7.5μl |
| SARS-CoV-2_20_RIGHT | AACACYTAAAGCAGCGGTTGAG | REV | 7.5μl |
| SARS-CoV-2_22_LEFT | AGTTGCAGAGTGGTTTTTGGCA | FWD | 10μl |
| SARS-CoV-2_22_RIGHT | ACTGTAGTGACAAGTCTCTCGCA | REV | 10μl |
| SARS-CoV-2_24_LEFT | AGCTAATAACACTAAAGGTTCATTGCCT | FWD | 10μl |
| SARS-CoV-2_24_RIGHT | TGACTTTTTGCTACCTGCGCAT | REV | 10μl |
| SARS-CoV-2_26_LEFT | TGGTTGAAGCAGTTAATTAAAGTTACACT | FWD | 10μl |
| SARS-CoV-2_26_RIGHT | TTCAGCAGCCAAAACACAAGCT | REV | 10μl |
| SARS-CoV-2_28_LEFT | CCTTGAAGGTTCTGTTAGAGTGGT | FWD | 10μl |
| SARS-CoV-2_28_RIGHT | AGGTGTGAACATAACCATCCACTG | REV | 10μl |
| SARS-CoV-2_30_LEFT | AGAAATGTATCTAAAGTTGCGTAGTGATG | FWD | 10μl |
| SARS-CoV-2_30_RIGHT | CCCTGAGTTGAACATTACCAGCC | REV | 10μl |
| SARS-CoV-2_32_LEFT | TACCAATGTGCTATGAGGCCCA | FWD | 10μl |
| SARS-CoV-2_32_RIGHT | GCACTACCCAATATGGTACGTCC | REV | 10μl |
| SARS-CoV-2_34_LEFT | GTCCAGAGTACTCAATGGTCTTTGT | FWD | 10μl |
| SARS-CoV-2_34_RIGHT | ACCTCTGGCCAAAAACATGACA | REV | 10μl |
| SARS-CoV-2_36_LEFT | CGCTACTTTAGACTGACTCTTGGTG | FWD | 10μl |
| SARS-CoV-2_36_RIGHT | ATCACCATTAGCAACAGCCTGC | REV | 10μl |
| SARS-CoV-2_38_LEFT | AGATCTGAGGACAAGAGGGCAA | FWD | 10μl |
| SARS-CoV-2_38_RIGHT | TGTCATCAGTGCAAGCAGTTTGT | REV | 10μl |
| SARS-CoV-2_40_LEFT | GGTATGGTACTTGGTAGTTTAGCTGC | FWD | 10μl |
| SARS-CoV-2_40_RIGHT | ACGATTGTGCATCAGCTGACTG | REV | 10μl |
| SARS-CoV-2_42_LEFT | TCTCTAACTACCAACATGAAGAAACAATTT | FWD | 20μl |
| SARS-CoV-2_42_RIGHT | GCAGTTAAAGCCCTGGTCAAGG | REV | 20μl |
| SARS-CoV-2_44_LEFT | TGGACCACTAGTGAGAAAAATATTTGTTG | FWD | 10μl |
| SARS-CoV-2_44_RIGHT | ACAGCCACCATCGTAACAATCA | REV | 10μl |
| SARS-CoV-2_46_LEFT | TGCAAAGAATAGAGCTCGCACC | FWD | 10μl |
| SARS-CoV-2_46_RIGHT | TGCATTAACATTGGCCGTGACA | REV | 10μl |
| SARS-CoV-2_48_LEFT | CTCTCTGACGATGCTGTTGTGT | FWD | 10μl |
| SARS-CoV-2_48_RIGHT | TGCGGTGTGTACATAGCCTCAT | REV | 10μl |
| SARS-CoV-2_50_LEFT | AGGAGGTATGAGCTATTATTGTAAATCACA | FWD | 10μl |
| SARS-CoV-2_50_RIGHT | GTTGTACCTCGGTAAACAACAGCA | REV | 10μl |
| SARS-CoV-2_52_LEFT | TGCAAATTATCAAAAGGTTGGTATGCA | FWD | 10μl |
| SARS-CoV-2_52_RIGHT | CCGAGGAACATGTCTGGACCTA | REV | 10μl |
| SARS-CoV-2_54_LEFT | TGGAGAAAAGCTGTCTTTATTTCACCT | FWD | 10μl |
| SARS-CoV-2_54_RIGHT | GCTTCTTCGCGGGTGATAAACA | REV | 10μl |
| SARS-CoV-2_56_LEFT | ACCACCGCCTGGAGATCAATTT | FWD | 10μl |
| SARS-CoV-2_56_RIGHT | CGCTTAACAAAGCACTCGTGGA | REV | 10μl |
| SARS-CoV-2_58_LEFT | GCCTTGTAGTGACAAAGCTTATAAAATAGA | FWD | 10μl |
| SARS-CoV-2_58_RIGHT | AAACCCACAAGCTAAAGCCAGC | REV | 10μl |
| SARS-CoV-2_60_LEFT | CAGGGTGAAGTACCAGTTTCTATCATT | FWD | 10μl |
| SARS-CoV-2_60_RIGHT | GAGTAAAGTAAGTTTCAGGTAATTGTTGG | REV | 10μl |
| SARS-CoV-2_62_LEFT | TCGTTTATGGAGATTTTAGTCATAGTCAGT | FWD | 10μl |
| SARS-CoV-2_62_RIGHT | TTGCGACATTCATCATTATGCCTTT | REV | 10μl |
| SARS-CoV-2_64_LEFT | CTGTACATACAGCTAATAAATGGGATCTCA | FWD | 10μl |
| SARS-CoV-2_64_RIGHT | TTTGACCTTCTTTTAAAGACATAACAGCA | REV | 10μl |
| SARS-CoV-2_66_LEFT | ACCCCCTGCATACACTAATTCTYT | FWD | 10μl |
| SARS-CoV-2_66_RIGHT | ACCCTGTTTTCCTTCAAGGTCC | REV | 10μl |
| SARS-CoV-2_68_LEFT | ACATAGAAGTTATTTGACTCCTGGTGA | FWD | 10μl |
| SARS-CoV-2_68_RIGHT | CCCTGGAGCGATTTGTCTGACT | REV | 10μl |
| SARS-CoV-2_70_LEFT | CCGGTAGCACACCTTGTAATGG | FWD | 10μl |
| SARS-CoV-2_70_RIGHT | CCCCTATTAAACAGCCTGCACG | REV | 10μl |
| SARS-CoV-2_72_LEFT | TGTTACCACAGAAATTCTACCAGTGT | FWD | 10μl |
| SARS-CoV-2_72_RIGHT | TACCCGCTAACAGTGCAGAAGT | REV | 10μl |
| SARS-CoV-2_74_LEFT | GTGCACTTGGAAAACTTCAAGATGT | FWD | 10μl |
| SARS-CoV-2_74_RIGHT | TGTTACAAACCAGTGTGTGCCA | REV | 10μl |
| SARS-CoV-2_76_LEFT | GTTGATTTAGGTGACATCTCTGGCA | FWD | 7.5μl |
| SARS-CoV-2_76_RIGHT | AGCGCTCTGAAAAACAGCAAGA | REV | 7.5μl |
| SARS-CoV-2_78_LEFT | CTTTGGCTTTGCTGGAAATGCC | FWD | 10μl |
| SARS-CoV-2_78_RIGHT | GTGCTTACAAAGGCACGCTAGT | REV | 10μl |
| SARS-CoV-2_80_LEFT | ACGTGAGTCTTGTAAAACCTTCTTTTT | FWD | 10μl |
| SARS-CoV-2_80_RIGHT | AATGACCACATGGAACGCGTAC | REV | 10μl |
| SARS-CoV-2_82_LEFT | GGACCTGCCTAAAGAAATCACTGT | FWD | 10μl |
| SARS-CoV-2_82_RIGHT | TGCCCTCGTATGTTCCAGAAGA | REV | 10μl |
| SARS-CoV-2_84_LEFT | CTTCACACTCAAAAGAAAGACAGAATGA | FWD | 10μl |
| SARS-CoV-2_84_RIGHT | ACGAACAACGCACTACAAGACT | REV | 10μl |
| SARS-CoV-2_86_LEFT | GGCCCCAAGGTTTACCCAATAA | FWD | 10μl |
| SARS-CoV-2_86_RIGHT | CTGTTGCGACTACGTGATGAGG | REV | 10μl |
| SARS-CoV-2_88_LEFT | TAACACAAGCTTTCGGCAGACG | FWD | 10μl |
| SARS-CoV-2_88_RIGHT | GTGGTCTGCATGAGTTTAGGCC | REV | 10μl |
